# Neuronal Network Dynamics in the Posterodorsal Amygdala: Shaping Reproductive Hormone Pulsatility

**DOI:** 10.1101/2024.01.21.574304

**Authors:** Kateryna Nechyporenko, Margaritis Voliotis, Xiao Feng Li, Owen Hollings, Deyana Ivanova, Jamie J Walker, Kevin T O’Byrne, Krasimira Tsaneva-Atanasova

## Abstract

Normal reproductive function and fertility rely on the rhythmic secretion of gonadotropin-releasing hormone (GnRH), which is driven by the hypothalamic GnRH pulse generator. A key regulator of the GnRH pulse generator is the posterodorsal subnucleus of the medial amygdala (MePD), a brain region which is involved in processing external environmental cues, including the effect of stress. However, the neuronal pathways enabling the dynamic, stress-triggered modulation of GnRH secretion remain largely unknown. Here, we employ in-silico modelling in order to explore the impact of dynamic inputs on GnRH pulse generator activity. We introduce and analyse a mathematical model representing MePD neuronal circuits composed of GABAergic and glutamatergic neuronal populations, integrating it with our GnRH pulse generator model. Our analysis dissects the influence of excitatory and inhibitory MePD projections’ outputs on the GnRH pulse generator’s activity and reveals a functionally relevant MePD glutamatergic projection to the GnRH pulse generator, which we probe with in vivo optogenetics. Our study sheds light on how MePD neuronal dynamics affect the GnRH pulse generator activity, and offers insights into stress-related dysregulation.

## 1. Introduction

The rhythmic secretion of gonadotropin-releasing hormone (GnRH) from the hypothalamus into the portal circulation is crucial in triggering the pulsatile release of gonadotropin hormones (luteinizing hormone (LH) and follicle-stimulating hormone (FSH)) from the pituitary gland [1, 2]. This dynamic process significantly contributes to the initiation of puberty and plays a pivotal role in ensuring fertility [2, 3]. The pulsatile release of GnRH is controlled by an upstream brain network known as the “GnRH pulse generator”. This network is composed of KNDy neurons found in the arcuate nucleus (ARC) of the hypothalamus, which co-express kisspeptin, neurokinin B and dynorphin A [4, 5]. We have previously used a mathematical model of the KNDy network to show that periodic pulsatile activity emerges as the basal activity or external activation of the network is increased, and confirmed this modelling prediction in vivo using optogenetic stimulation of the KNDy network [6, 7].

The GnRH pulse generator is sensitive to inputs from neural centres relaying information about the psychological and physiological state of the organism. In particular, the posterodorsal subnuclei of the medial amygdala (MePD), a limbic structure responsible for emotional processing of complex external cues, is responsive to stress [8] and regulates pubertal onset [9–11] as well as LH pulsatility [12, 13]. Whilst there is limited understanding of how the MePD processes stress-related information and relays it to the GnRH pulse generator, it has been shown that optogenetic stimulation of kisspeptin neurons in the MePD increases LH pulse frequency [12]. This effect is mediated by both GABAergic and glutamatergic signalling within the MePD, since pharmacological antagonism of GABA receptors within the MePD prevents the increase in LH pulse frequency, while pharmacological antagonism of glutamate receptors within the MePD terminates the LH pulses altogether [13]. It is likely that MePD regulation of LH pulsatility is mediated, at least in part, through direct projections from the MePD to KNDy neurons within the ARC. Indeed, viral-based monosynaptic tract-tracing in mice has shown that the amygdala provides inputs to ARC KNDy neurons [14, 15]. Furthermore, although glutamatergic projections from the MePD have not yet been explored experimentally, it has been shown that stimulation of MePD GABAergic projection terminals in the ARC causes a suppression of LH pulses [16].

Taken together, the available data suggest that the MePD can modulate GnRH pulse generator activity through local kisspeptin signalling. However, how the GABA-glutamate neuronal network within the MePD integrates kisspeptin activity and relays this to the GnRH pulse generator is less clear. Here, we investigate this using a mathematical mean-field model of the MePD GABA-glutamate neuronal circuit (see Figure 1(**a**)). We calibrate the model using available experimental observations and analyse it to understand how changes in functional connectivity within the MePD’s neuronal network affect the system’s output. We then couple this model of the MePD with our earlier mathematical model of the GnRH pulse generator [6, 7] to study the effects of manipulating both GABAergic and glutamatergic signalling on the activity of the GnRH pulse generator.

**Figure 1.**
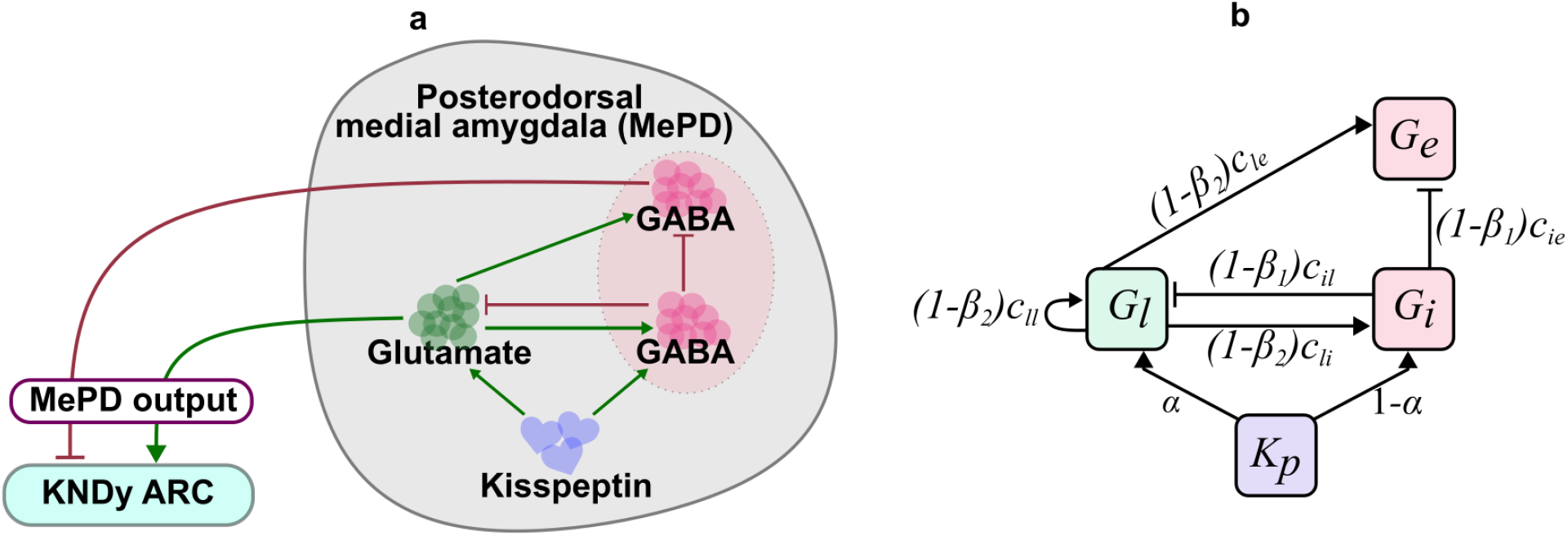
(**a**) Schematic diagram of the GABA-glutamate circuit in the MePD. Kisspeptin activates glutamatergic and GABAergic neuronal populations in the MePD. Both glutamatergic and GABAergic populations interact functionally with a population of GABA efferent neurons providing excitation and inhibition, respectively. The activity within the GABAergic efferent neurons and glutamatergic neurons extends to the ARC to modulate the activity of the KNDy population. (**b**) Network representation of the MePD circuit model depicting interactions between the populations of glutamatergic neurons(*G*_*l*_), GABA interneurons (*G*_*i*_) and GABAergic efferents (*G*_*e*_). Both *G*_*l*_ and *G*_*i*_ receive kisspeptin excitation (*K*_*p*_), that is distributed proportionally with relative glutamatergic excitation ratio *α*. The parameters *c* represent the strength of functional interactions between the populations. Parameters *β*_1_ and *β*_2_ are the interaction suppression coefficients, used to mimic the decrease in the GABAergic and glutamatergic interaction strengths, respectively, in order to simulate the effects of chemical antagonism.

## 2. Methods

### (a) Animals

Adult Vglut-flp mice heterozygous for the allele Slc17a6<tm1.1(flpo)Hze> (Strain #:030212, B6;129S-S*l*c17a6tm1.1(flpo)Hze*/*J; Jackson Laboratory, Bar Harbor, ME, USA) were bred in house. Mice were genotyped by PCR using the following primers: mutant reverse, 13007-ACA CCG GCC TTA TTC CAA G; common, 34763-GAA ACG GGG GAC ATC ACT C; and wildtype reverse, 34764-GGA ATC TCA TGG TCT GTT TTG. Mice, aged 7-9 weeks at time of initial surgery, were group housed unless chronically implanted with fiber optic cannulae and kept at 25°C ± 1°C, 12:12 hr light/dark cycle (lights on 0700 h), with ad libitum access to food and water. All procedures were performed in accordance with UK home office regulation and approved by The King’s College London Animal Welfare and Ethical Review Body.

### (b) Stereotaxic injection of viral constructs and implantation of fibre optic cannula

All surgical procedures were carried out under general anaesthesia using ketamine (Vetalar, 100 mg/kg, i.p.; Pfizer, Sandwich, UK) and xylazine (Rompun, 10 mg/kg, i.p.; Bayer, Leverkusen, Germany) under aseptic conditions. Mice were secured in a David Kopf stereotaxic frame (Model 900, Kopf Instruments) and bilaterally ovariectomised (OVX). A midline incision of the scalp was used to expose the skull. The periosteum was removed and two small bone screws were inserted into the skull. Using a robot stereotaxic system (Neurostar, Tubingen, Germany) two windows were drilled intracranially directly above target coordinates for the posterodorsal subnucleus of the medial amygdala (MePD) (2.35 mm lateral, 1.45 mm posterior to bregma, at a depth of 5.49 mm below the skull surface) and arcuate nucleus (ARC) (0.24 mm lateral, 1.51 mm posterior to bregma, at a depth of 5.85 mm below the skull surface) obtained from the mouse brain atlas of [17]. Either AAV-CAG-FLEXFRT-ChR2(H134R)-mCherry (*n* = 7, 200 nL, 3 × 1012 GC*/*mL, Serotype: 9; Addgene, MA, USA) to express channelrhodopsin (ChR2) or control virus (*n* = 4, AAV-Ef1a-fDIO-mCherry, 200 mL, 3 × 1012 GC/mL; Serotype: 9; Addgene) was unilaterally injected over 10 minutes into the MePD using a 2-*µ*L Hamilton microsyringe (Esslab, Essex, UK). The needle was left in place for 5 minutes, before withdrawing 0.2 mm, waiting another minute, and then fully withdrawing over the course of 1 minute. A fiber optic cannula (200 *µ*m,0.39 NA, 1.25 mm ceramic ferrule, Doric Lenses, Quebec, Canada) was inserted into the brain targeting the ARC. This was fixed to the skull surface and bone screws using dental cement (Super-Bond Universal Kit, Prestige Dental, UK) before suturing closed the incision. A one week recovery period was given post-surgery. After this period, the mice were handles daily to acclimatize them to the tail-top blood sampling procedure [18]. Mice were left for 4 weeks to achieve effective opsin expression before experimentation.

### (c) Experimental design and blood sampling for LH measurement

For measurement of LH pulsatility during optogenetic stimulation, the tip of the mouse’s tail was removed with a sterile scalpel for tail-tip blood sampling [19]. The chronically implanted fiber-optic cannula was attached to a multimode fiber-optic rotary joint patch cables (Thorlabs, Ltd, Ely, UK) via a ceramic mating sleeve which allows mice to freely move while receiving blue light (473 nm wavelength). Laser (DPSS laser, Laserglow Technologies, Toronto, Canada) intensity was set to 10 mW at the tip of the fibre optic patch cable. The frequency and pattern of optical stimulation was controlled by software designed in house. After 1 h acclimatization, blood samples (5*µ*l) were collected every 5 min for 2 h. After 1 h controlled blood sampling without optical stimulation to determine baseline LH pulse frequency, mice received patterned optical stimulation (5 s on 5s off, 10-ms pulse width) at 2, 5, 10 or 20 Hz for 1 h while blood sampling continued. Non-stimulation controls were performed in the same manner, with no stimulation during the second hour. The control virus injected animals received 5 Hz optical stimulation. Mice received all treatments in a random order, with at least 3 but typically 5 days between experiments.

The blood samples were snap-frozen on dry ice and storing at −80°C until processed. In-house LH enzyme-linked immunosorbent assay (LH ELISA) similar to that described by [18] was used for processing of the mouse blood samples. The mouse LH standard (AFP-5306A; NIDDK-NHPP) was purchased from Harbor-UCLA along with the primary antibody (polyclonal antibody, rabbit LH antiserum, AFP240580Rb; NIDDK-NHPP). The secondary antibody (donkey anti-rabbit IgG polyclonal antibody [horseradish peroxidase]; NA934) was from VWR International. Validation of the LH ELISA was done in accordance with the procedure described in [18] derived from protocols defined by the International Union of Pure and Applied Chemistry. Serially diluted mLH standard replicates were used to determine the linear detection range. Nonlinear regression analysis was performed using serially diluted mLH standards of known concentration to create a standard curve for interpolating the LH concentration in whole blood samples, as described previously [6]. The assay sensitivity was 0.031 ng/mL, with intra- and inter-assay coefficients of variation of 4.6% and 10.2% respectively.

### (d) LH pulse detection and statistical analysis

LH pulses were determined using the DynPeak algorithm [20], with settings adjusted to accommodate for the high LH pulse frequency in OVX mice as previously outlined by Breen and colleagues [21]. These include using the programmes’ default parameters except the global threshold was increased to 35%, the nominal peak threshold was reduced to 20 min and the 3-point peak threshold was removed. Average LH inter-pulse interval (IPI) (the period of time between two LH pulse peaks) was calculated for the 1 h control period and 1 h optogenetic stimulation period or equivalent non-stimulation period in control animals. Statistical significance was tested using a two-way repeated measures ANOVA and post-hoc Tukey test. Data was represented as mean ± SEM and *p <* 0.05 was considered significant.

### (e) Mean-Field Model of the MePD

Given the established presence of GABA and glutamate neuronal populations in the MePD [22, 23], we model their interplay employing Wilson-Cowan framework [24, 25]. The framework allows us to take a system-level approach to describe the dynamic evolution of excitatory/inhibitory activity in neuronal populations due to functional interactions within a synaptically-coupled neuronal network, incorporating both cooperation and competition mechanisms. Rather than considering individual neurons within the populations, the framework gives a coarse-grained account of the mean activity in the network, which allows the investigation of putative functional interactions between the various populations as well as the overall network output, enabling coupling of our MePD network model to other neuronal networks, such as the GnRH pulse generator as represented by the KNDy network in the ARC[6, 7].

The Wilson-Cowan framework includes an inhibitory and excitatory dependant variables, both receiving an excitatory input. However, straightforward application of this framework is not sufficient to describe the MePD GABA-glutamate neuronal network to represent differential dynamics of GABA-mediated disinhibitory mechanism [26, 27]. Therefore, we have extended the original Wilson-Cowan model by incorporating an additional inhibitory population (GABA), that does not receive excitatory input (see Figure 1(**b**)).

A key component of the Wilson-Cowan modelling framework is a sigmoid stimulus-response function *ϕ*(*a, F, θ*), which controls the mean level of activity generated in the populations at a time *t* [24]:

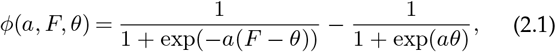

where *F* indicates the input to a given population and parameters *a* and *θ* define the value of maximum slope and half-maximum firing threshold, respectively. The use of the sigmoidal function is motivated by the fact that the majority of neurons have fluctuating m embrane p otential n ear an excitability threshold, such that the probability of firing grows exponentially upon depolarisation [28]. Additionally, the constant 1*/*(1 + exp(*aθ*)) ensures that under the absence of stimulatory input to the population the firing ceases, i.e. *ϕ*(*a*, 0, *θ*) = 0 [24].

The input *F* is given by the linear sum of excitatory and inhibitory contributions, as follows:

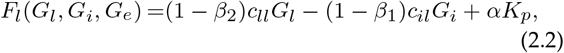

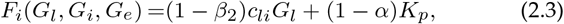

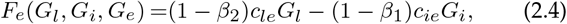

where the parameters *c* represent the strength of interaction from one population to another, as shown in Figure 1(**b**). The parameter *K*_*p*_ depicts the overall kisspeptin level of excitatory input to the system, which is then distributed to the populations of glutamatergic neurons and GABA interneurons in accordance with the relative glutamatergic excitation ratio parameter *α* ∈ [0, 1] representing the proportion of input directed to glutamatergic neuronal population. We minimised the number of inhibitory coupling strength parameters by setting self-inhibition in the GABAergic populations and functional interactions between GABAergic efferent neurons and the other two populations to zero. In the absence of data that specifically supports the inclusion of such inhibitory interactions, a model with fewer parameters is justified and easier to interpret. As one of our aims is to investigate effects of GABA and glutamate receptor antagonism following pharamcological interventions, we incorporate the terms (1 − *β*_1_) and (1 − *β*_2_), where *β*_1_ and *β*_2_ represent the proportion of suppressed functional interaction between GABA and glutamate neuronal populations, respectively.

Using the stimulus-response function and the proposed interactions between the populations (Figure 1(**b**)) the averaged activity in the populations is governed by the following functions:

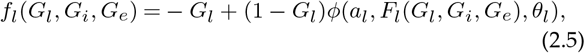

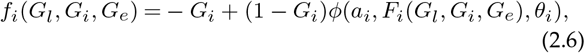

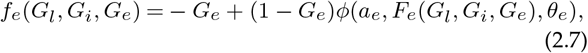

where the dependent variables *G*_*l*_, *G*_*i*_, *G*_*e*_ represent the mean activity in the populations of glutamatergic neurons, GABA interneurons and GABAergic efferent neurons at time *t*, respectively. The model also includes refractory dynamics via the term (1 − *G*), which controls the time period during which the populations are unable to produce a signal following an activation, and its primary effect is decreasing the maximum firing rate [29]. The MePD activity is then governed by the following ordinary differential equations (ODEs):

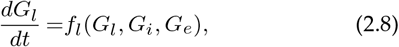

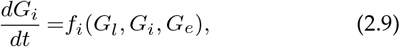

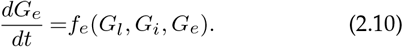

The presented MePD system is non-dimensional w.r.t. time [30]. To effectively couple the system, we introduce the time scaling factor *d* that relates arbitrary time in equations 2.8-2.10 to time in minutes:

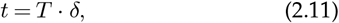

where *t* is the original arbitrary time, *T* is the new time measured in minutes, and *δ* is the scaling factor (min^*−*1^). The time-converted version of the model is as follows:

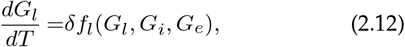

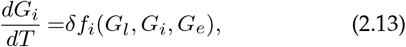

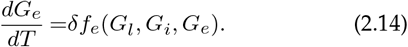

The parameter values for the MePD model can be found in Table 1.

**Table 1.** MePD model parameter values.

| Parameter | Definition | Value | Reference |
| --- | --- | --- | --- |
| $\delta$ | Temporal scaling factor ($\text{min}^{-1}$) | 3 | [31] |
| $\alpha$ | relative glutamatergic excitation ratio (a.u.) | 0.9 | Derived |
| $K_p$ | Excitatory input to the MePD circuit (a.u.) | [0, 14] | |
| $c_{ll}$ | glutamatergic self-excitation strength ($\text{Glu} \rightarrow \text{Glu}$) (a.u.) | 18 | Derived |
| $c_{li}$ | interaction strength $\text{Glu} \rightarrow \text{GABA}_{\text{int}}$ (a.u.) | 16 | Derived |
| $c_{il}$ | interaction strength $\text{GABA}_{\text{int}} \rightarrow \text{Glu}$ (a.u.) | 35 | Derived |
| $c_{le}$ | glutamatergic excitation of GABA efferents ($\text{Glu} \rightarrow \text{GABA}_{\text{eff}}$) (a.u.) | 40 | Derived |
| $c_{ie}$ | GABAergic inhibition of GABA efferent neurons ($\text{GABA}_{\text{int}} \rightarrow \text{GABA}_{\text{eff}}$) (a.u.) | 25 | Derived |
| $a_l$ | maximum slope of $\text{Glu}$ (a.u.) | 1.3 | [24] |
| $a_i$ | maximum slope of $\text{GABA}_{\text{int}}$ (a.u.) | 2 | [24] |
| $a_e$ | maximum slope of $\text{GABA}_{\text{eff}}$ (a.u.) | 2 | [24] |
| $\theta_l$ | half-maximum firing threshold for $\text{Glu}$ (a.u.) | 4 | [24] |
| $\theta_i$ | half-maximum firing threshold for $\text{GABA}_{\text{int}}$ (a.u.) | 3.7 | [24] |
| $\theta_e$ | half-maximum firing threshold for $\text{GABA}_{\text{eff}}$ (a.u.) | 3.7 | [24] |
| $\beta_1$ | GABAergic interaction suppression coefficient (a.u.) | [0, 1] | |
| $\beta_2$ | glutamatergic interaction suppression coefficient (a.u.) | [0, 1] | |

### (f) Calculating MePD output in the mean-field model

The magnitude of the mean glutamatergic and GABAergic MePD projections’ output is found in the same way, i.e. as the integral of mean activity in the respective population over the integration time period *T* :

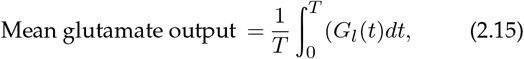

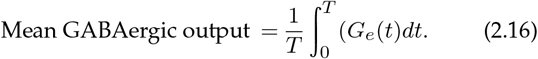

To quantify the periodic output of the MePD GABA-glutamate circuit we compute the integral of the difference of mean activity in the populations of glutamatergic neurons (*G*_*l*_) and GABAergic efferent neurons (*G*_*e*_) over the integration time period *T* :

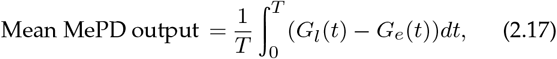

where the term 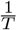 is the reciprocal of the time duration, allowing to normalise the output.

### (g) Coarse-grained Model of ARC KNDy population with MePD input

Based on experimental evidence regarding MePD projections to other brain regions including the ARC [14, 15] we couple our MePD model’s output with our ARC KNDy network model [6, 7] aiming to explore the effects ofperturbations to the MePD circuit on GnRH pulse generator activity and to validate our model against experimental observations in [12, 13]. The model describing the dynamics in the KNDy neuronal network is given by the following system of ordinary differential equations:

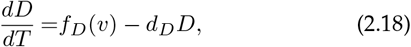

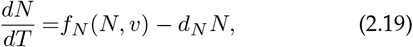

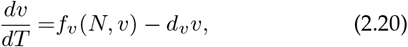

where *D* and *N* represent the concentration of Dynorphin and Neurokinin B produced by the population, and *v* describes the averaged firing activity in the population in spikes/min. Parameters *d*_*D*_,*d*_*N*_, and *d*_*v*_ control the linear decay for each variable. Dynorphin and Neurokinin B secretion rates are represented by functions *f*_*D*_ and *f*_*N*_, respectively, while *f*_*v*_ describes how the firing rate changes in response to the Neurokinin B concentration and current firing rate. The neuropeptides’ secretion rates are given by the following functions:

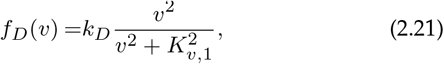

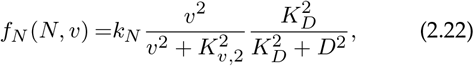

where *k*_*D*_ and *k*_*N*_ signify the neuropeptides’ secretion rates; *K*_*v*,1_ and *K*_*v*,2_ describe the frequency value for which the rate of Dynorphin and Neurokinin B secretion is half-maximum; and *K*_*D*_ describes the Dynorphin concentration that results in half-maximum inhibition of NKB. In the original introduction of the KNDy model [6], the function *f*_*v*_ can take both positive and negative values. Here we modified *f*_*v*_ by restricting its output to be non-negative via vertical shift of the sigmoid function:

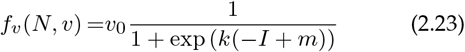

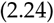

where *v*_0_ is the maximum increase of the firing rate in response to synaptic inputs *I* (Hz). The parameter *m* signifies the synaptic input level at which the increase in the firing rate becomes half-maximum. The parameter *k* represents the membrane’s time constant, which determines how quickly the neuron’s membrane potential changes in response to inputs.

The synaptic inputs *I* consist of both external and internal contributions:

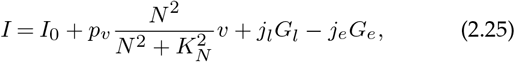

where *I*_0_ stands for the basal input in the population. The excitatory effect of Neurokinin B on the firing rate is accounted via a sigmoid function with *K*_*N*_ representing Neurokinin B’s half-maximal effect and *p*_*v*_ control’s the strength of the connection between the neurons in the KNDy. To account for the effects of the MePD output on the KNDy, we use the terms *j*_*l*_*G*_*l*_ and −*j*_*e*_*G*_*e*_, which signify the excitatory glutamatergic and inhibitory GABAergic contribution from the MePD, respectively. The parameters *j*_*l*_ and *j*_*e*_ are presynaptic firing rate conversion parameters for the corresponding populations in the MePD. These parameters allow converting non-dimensional output from the MePD to the input to the KNDy in Hz and assign weight to the contribution and, we alter when simulating the effects of MePD projections’ stimulation. In the case of simulating the effects of neurotransmitter antagonism and kisspeptin stimulation in the MePD we set *j*_*l*_ = *j*_*e*_ = 1. To mimic the effects of stimulating glutamatergic and GABAergic projections, we consider varied levels of MePD network excitation (*K*_*p*_) and alter the presynaptic firing rate conversion parameters to change the weight distribution of the projections. In case of glutamatergic projections stimulation we increase the weight of the glutamatergic contribution and decrease the GABAergic weight contribution (*j*_*l*_ = 1.5, *j*_*e*_ = 0.5), while for GABA projections we increase the weight of the GABAergic contribution and decrease the glutamatergic weight contribution (*j*_*l*_ = 0.5, *j*_*e*_ = 1.5). For further details on the original KNDy model, refer to [6]. The parameters for the KNDy network can be found in Table 2.

**Table 2.** KNDy model parameter values.

| Parameter | Definition | Value | Reference |
| --- | --- | --- | --- |
| $d_D$ | Dyn degradation rate ($\text{min}^{-1}$) | 0.2 | [6] |
| $d_N$ | NKB degradation rate ($\text{min}^{-1}$) | 1 | [6] |
| $d_v$ | Firing rate reset rate ($\text{min}^{-1}$) | 10 | [32] |
| $k_D$ | Dyn signalling strength ($\text{nMmin}^{-1}$) | 4 | [7] |
| $k_N$ | NKB signalling strength ($\text{nMmin}^{-1}$) | 40 | [7] |
| $p_v$ | Effective strength of synaptic input (a.u.) | 0.006 | [7] |
| $v_0$ | Maximum rate of neuronal activity increase ($\text{spikes min}^{-2}$) | 25000 | [32] |
| $K_D$ | Dyn $\text{IC}_{50}$ (nM) | 0.3 | [33] |
| $K_N$ | NKB $\text{IC}_{50}$ (nM) | 4 | [34] |
| $K_{v,1}$ | Firing rate for half-maximal Dyn secretion ($\text{spikes min}^{-1}$) | 600 | [35] |
| $K_{v,2}$ | Firing rate for half-maximal NKB secretion ($\text{spikes min}^{-1}$) | 200 | [35] |
| $k$ | Membrane's time constant (min) | 10 | Fixed |
| $m$ | Half-maximal firing rate ($\text{min}^{-1}$) | 0.5 | Fixed |
| $I_0$ | Basal activity ($\text{min}^{-1}$) | 0.14 | Fixed |
| $j_l$ | presynaptic firing rate conversion parameter for glutamatergic projections (Hz) | {1, 1.5, 0.5} | |
| $j_e$ | presynaptic firing rate conversion parameter for GABAergic projections (Hz) | {1, 0.5, 1.5} | |

### (h) Numerical simulations and bifurcation analysis

Bifurcation analysis was performed in AUTO 07-p [36], while numerical simulations were carried out in MATLAB using ode45 (Runge-Kutta method) for the MePD system and ode15s (variable-step, variable-order (VSVO) solver) for the coupled MePD-KNDy model. The codes for reproducing the analysis and simulations presented in this manuscript can be found in [GitHub repository](https://github.com/ktrnnchprnk/MePDmodel).

## 3. Results

### (a) Modelling the MePD’s GABA-glutamate circuit and its projections to the ARC

The diagram depicted in Figure 1(**a**) provides an overview of our model describing the functional connectivity in the MePD neural circuit along with the MePD’s projections (outputs) to the ARC. In our mode we consider an excitatory population of glutamatergic neurons and an inhibitory population of GABAergic neurons given the experimentally-established presence of glutamatergic and GABAergic neurons in the MePD [22, 23]. These populations of glutamatergic and GABAergic neurons interact with each other, and also extend excitatory and inhibitory connections, respectively, to a distinct neuronal population of GABAergic neurons [26, 27], which we refer to here as GABA efferent neurons. The activity in the populations of the GABA efferent neurons and glutamatergic neurons defines the MePD’s output that we consider in the model to be acting on the ARC. It has previously been shown by [13] that a kisspeptin-expressing neuronal population, found in the MePD [37], provides excitatory input to the populations of GABA interneurons and glutamatergic neurons and has been accordingly included in our modelling. Our mathematical model is based on the established Wilson-Cowan modelling framework [24, 25] and hence allows us to simulate the mean activity of the different neuronal populations; namely glutamatergic neurons (Glu), GABA interneurons (GABA_int_) and GABA efferent neurons (GABA_eff_).

In this study, we couple the MePD neuronal network model to our KNDy neuronal network model [6, 7]. In previous work, we coupled a first-generation model of the MePD neuronal network with our KNDy (pulse generator) model and performed numerical simulations in order to reproduce the results of optogenetic stimulation of kisspeptin and pharmacological antagonism experiments in the MePD in [13]. The MePD neuronal network model used in [13] was based on the same framework as the model in this manuscript, but under the assumption of stationary MePD network activity and hence constant MePD output. In the present study, we investigate the proposed MePD neuronal network in more detail, taking into account the possibility of dynamic (e.g. oscillatory) MePD activity. To this end, we match the temporal activity in the circuit to the time scales of calcium activity recorded in MePD neurons [31]. Such activity is now routinely used as a proxy of mean neuronal activity, and in our case it is mediated by GABA and glutamate neuronal populations in the MePD. Full details of the model are given in Mean-Field Model of the MePD.

### (b) How does excitatory input decrease inhibitory tone in the MePD circuit?

Optogenetic stimulation of MePD kisspeptin neurons has been shown to have a significant effect on LH pulses [12], presumably via exciting GABA-glutamate neuronal circuits and their projections to the ARC. To investigate how this effect could be relayed through the GABA-glutamate MePD neuronal network, we study the model’s behaviour under various levels of excitatory kisspeptin input. Previous analysis of the Wilson-Cowan model [24, 25, 30] has shown that oscillatory dynamics in the model can be induced via glutamatergic self-excitation and a negative-feedback loop between the populations of GABA interneurons and glutamate neurons. Accordingly, in our MePD model, we assume excitatory coupling from the glutamatergic population to the population of GABA interneurons as well as inhibitory coupling from the GABA interneurons to the glutamatergic population. To mimic experimental optogenetic stimulation of the GABA-glutamate network, we perform a bifurcation analysis using the level of MePD kisspeptin (“MePD excitation” in Figure 2) as a free parameter (for further details on the numerical methods, see Numerical simulations and bifurcation analysis).

**Figure 2.**
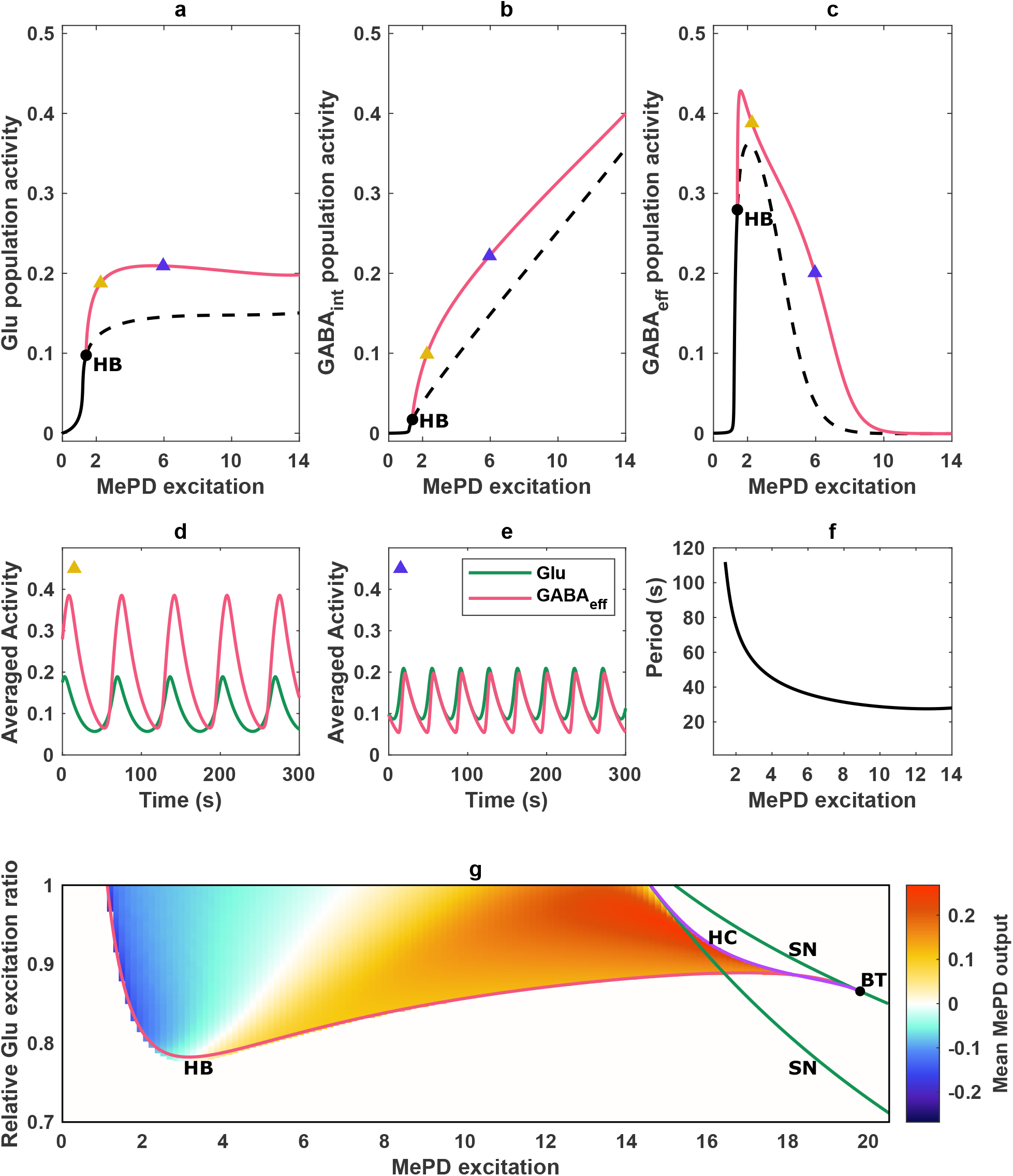
MePD GABA-glutamate circuit dynamics. (**a**-**c**) One-parameter bifurcation diagrams of the model for varying kisspeptin (*K*_*p*_) excitation of the GABA-glutamate circuit (MePD excitation). The qualitative dynamics in the populations of (**a**) glutamatergic neurons (Glu), (**b**) GABA interneurons (GABA_int_) and (**c**) GABAergic efferent neurons (GABA_eff_) changes as the kisspeptin excitation level provided to the circuit varies. Circular markers denote a Hopf (**HB**) bifurcation that gives rise to the oscillatory behaviour in the model. The red line shows the maximum amplitude of the limit cycle solution branch. (**d**-**e**) Simulation of population activity of Glu (green) and GABA_eff_ (red) at two different levels of MePD excitation (*K*_*p*_ = 2.3 and *K*_*p*_ = 6), corresponding to the yellow and blue triangular markers in panels **a**-**c**. (**f**) MePD excitation vs period of the system’s limit cycle oscillations. (**g**) Two-parameter bifurcation diagram using the level of MePD excitation (*K*_*p*_) and the relative Glu excitation ratio (*α*) as free parameters. Superimposed heat map indicates the mean MePD output in the oscillatory region. Red, green, and purple lines represent Hopf (**HB**), saddle-node (**SN**) and homoclinic bifurcations (**HC**), respectively. Circular markers depict a Bogdanov-Takens (**BT**) point.

The analysis reveals that at low kisspeptin excitation, the activity of all neuronal populations considered in our model is low and exhibits stationary dynamics (Figure 2(**a-c**)). This could be explained by the fact that the system does not receive enough excitation to sustain oscillations, and as a result, settles in a state of low mean activity where each population behaves as a pool of independent single neuron oscillators. As excitation increases, the activity in all three populations is amplified. Numerical continuation along the stable equilibrium branch reveals that the system undergoes a change in qualitative dynamics (at *K*_*p*_ = 1.4) due to a Hopf bifurcation (**HB**), giving rise to a branch of stable periodic (limit cycle) solutions. Within the parameter range where the limit cycle solutions exist, at a threshold kisspeptin level of excitatory input (*K*_*p*_ = 1.6) the gain in the population of GABA efferent neurons switches from positive to negative (Figure 2(**c**)). As a result, further increase in excitation leads to decrease in the activity of the GABA efferent neuronal population, which approaches zero with further increase in the kisspeptin excitatory input (see Figure 2(**c**)). This can be explained by the fact that inhibitory input from GABA interneurons to GABA efferents outweighs the excitatory input from the glutamatergic population (Figure 2(**a-b**)). The presence of a negative feedback loop in the system leads to a higher rate of increase in activity of the GABA interneuron population compared to the population of glutamate neurons. Therefore, by comparing the activity of the populations of glutamatergic neurons and GABA efferents at different excitation levels (Figure 2(**d-e**)) we observe that, overall, the MePD projections’ output would increase due to the reduction in the inhibitory GABAergic tone. In the parameter range where the model exhibits periodic behaviour, we also find that the oscillatory period decreases as we increase excitation (Figure 2(**f**)). The numerical range of the oscillation period confirms that the temporal activity in the model aligns well with the experimentally observed average calcium oscillation period reported in [31], where the oscillations in MePD neuronal activity occur, approximately, in the span of a minute. We further extend the bifurcation diagram (see Figure S1), identifying that the further increase in excitation leads to an exponential increase in the oscillation period and subsequent destruction of the limit cycle solution via a global homoclinic bifurcation. Following further increase in the kisspeptin level, the system enters a bistable regime, induced and destroyed via saddle-node bifurcations, followed by constant a high population activity mode.

It is unknown whether MePD kisspeptin directly modulates GABA interneurons and/or glutamatergic neurons. To investigate the role of the distribution of excitatory input between the inhibitory (GABA) and excitatory (glutamate) populations in the model, we introduce a parameter that controls the relative kisspeptin excitation ratio (*α*). We are thus able to continue the loci of the co-dimension one Hopf and saddle-node bifurcations (electronic supplementary material, figure S1) in two-parameter space (namely, the excitatory input *K*_*p*_ and the relative kisspeptin excitation ratio *α*) (Figure 2(**g**)). The Hopf bifurcation and saddle-node bifurcation curves coalesce in the two-parameter space, where a co-dimension two Bogdanov-Takens (**BT**) point emerges. This (**BT**) point is also related to the appearance of a homoclinic bifurcation curve, representing a situation where the stable and unstable manifolds of a saddle equilibrium intersect, indicating the presence of complex dynamical behaviour in the system. The bifurcation curves representing the Hopf (**HB**), saddle node (**SN**) and homoclinic (**HC**) loci allow us to identify regions in two-parameter space characterised by different qualitative dynamics in the system; and in particular, regions where the system oscillates. We note that for the current choice of parameters (Table 1), the oscillations in the system can be induced only under the condition that the majority of excitation is directed to the excitatory population of glutamatergic neurons. To investigate how the distribution and different levels of excitation affect the output of the system within the oscillatory region, we compute a heat map depicting changes in the mean MePD projections’ output (Figure 2(**g**)) due to changes in activity of the populations of excitatory (glutamate) neurons and inhibitory (GABA) efferent neurons in our MePD model. For complete details on how the mean MePD projections’ output is defined and computed, see Calculating MePD output in the mean-field model. We find that an increase in the level of MePD kisspeptin excitation leads to a transition from inhibitory to excitatory MePD output. As the proportion of kisspeptin excitation to the glutamatergic population is increased, the inhibitory tone of the MePD circuit can be maintained under stronger excitation in the model (Figure 2(**g**)). This suggests that additional (kisspeptin) excitation of the glutamatergic population may lead to an increase in GABAergic tone, depending on the functional interaction strength between the glutamatergic and GABAergic neuronal populations. Taken together, our theoretical findings suggest that the reduction in GABA efferent neuron activity amid increased excitation of the MePD neuronal circuit may be reliant on the intricate interplay between competitive inhibitory and excitatory connections to the population of GABA efferent neurons in the MePD circuit.

### (c) How does MePD functional network connectivity affect MePD projections’ output?

The presence of GABAergic and glutamatergic neuronal populations in the MePD [22, 23], as well as their importance in the modulation of GnRH pulse generator activity, has been demonstrated experimentally [13]. However, the role of their functional interactions within the circuit remains unknown. Hence, in this section we investigate how changes in the functional interaction (coupling) strength affect the dynamics of the GABA-glutamate circuit and the MePD output. The aim here is to characterise network interaction patterns associated with oscillatory dynamics and the corresponding periodic MePD excitatory/inhibitory projections’ output.

The stimulatory effect of the amygdala on the GnRH pulse generator under the stimulation of kisspeptin neurons has been attributed to the activation of GABAergic interneurones, which in turn inhibit GABAergic efferent neurons [13], forming a GABA-GABA disinhibitory interaction, which is of interest in understanding functional mechanisms and dynamics in the MePD circuit. As competing excitatory and inhibitory signals counterbalance each other, we fix the interaction strength responsible for glutamate input to GABA efferent neurons and investigate how MePD projections’ output changes under the variation of kisspeptin (level) stimulation and the strength of interaction between GABA interneurons and GABA efferents (*c*_*ie*_) Figure 3. The mean glutamatergic projections’ output remains relatively unaffected by the strength of interaction due to the lack of GABA efferent neuronal input to the glutamate neuronal population, but its magnitude moderately increases as the excitation in the circuit increases (Figure 3(**a**)). Meanwhile, as the system is excited, a low strength of interaction results in an increase in the magnitude of the mean GABAergic projections’ output (Figure 3(**b**)), as the GABA-GABA interaction is not sufficient to induce a decrease in the activity of the population of GABA efferent neurons under the increased excitation in the MePD circuit. On the other hand, under high interaction strength, the population of GABA interneurons exerts an excessive inhibitory input to the population of GABA efferent neurons, resulting in a reduction in the GABA efferent neuronal population activity to zero (Figure 3(**b**)). We compute the combined MePD projections’ output as the difference in magnitude between the glutamatergic and the GABAergic projections’ outputs. We show that a low level of functional interaction strength between GABA interneurons and GABA efferents leads to a predominantly inhibitory mean MePD projections’ output, whereas a high interaction strength results in an excitatory mean MePD projection’s output that remains mostly unchanged under further excitation of the MePD circuit (Figure 3(**c**)). This happens as the change in the activity of the population of GABA efferent neurons is much higher compared to the population of glutamate neurons (Figure 3(**a-b**)). Hence, the mean MePD projections’ output is heavily dependent on the strength of GABA-GABA disinhibition.

**Figure 3.**
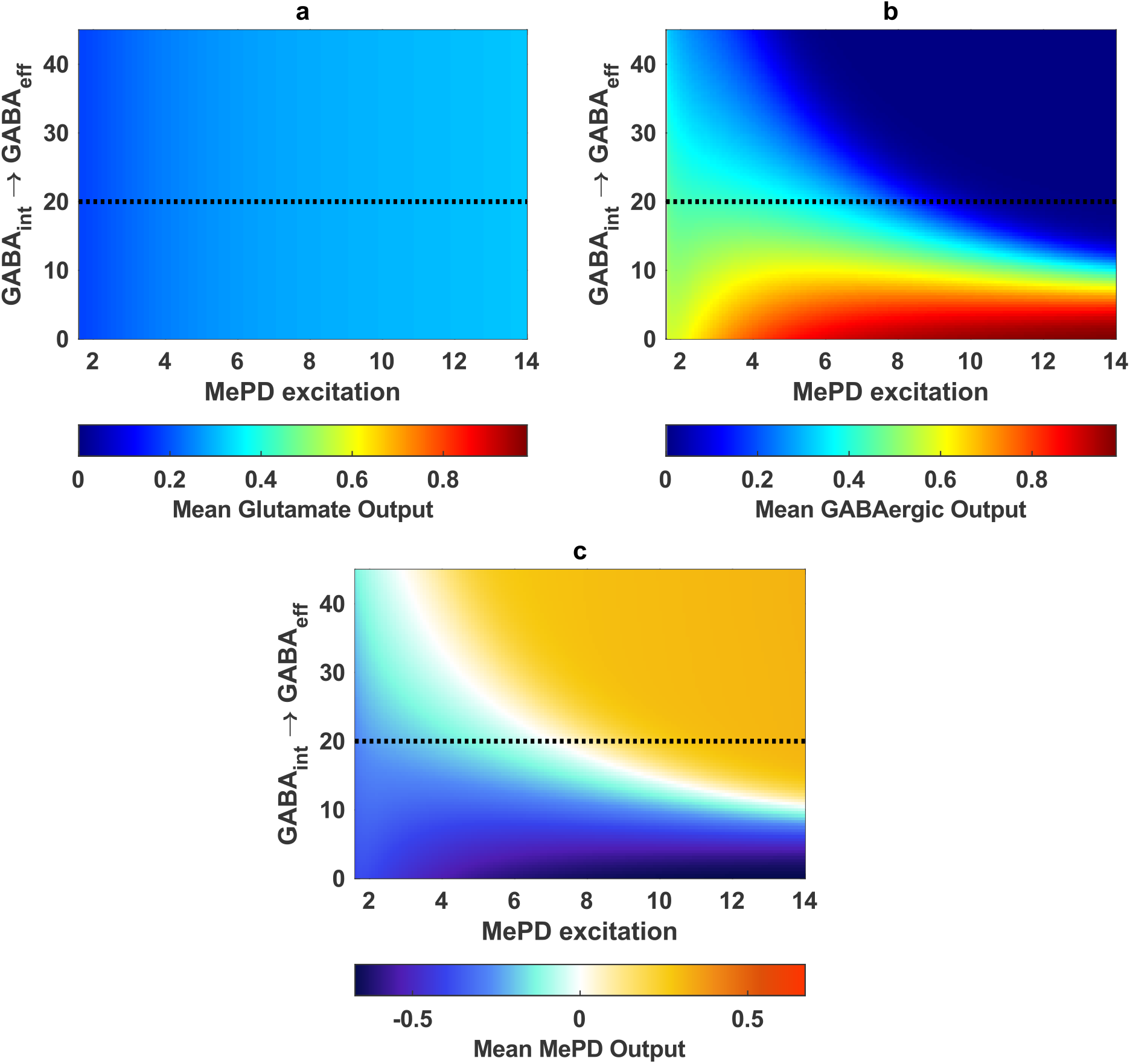
Effect of functional interaction strength between the population of GABA interneurons (GABA_int_) and the population of GABA efferent neurons GABA_eff_ on the MePD projections’ output. Heat map depicting (**a**) the magnitude of mean glutamatergic projections’ output, (**b**) the magnitude of the mean GABAergic output under varying MePD excitation (*K*_*p*_) and strength of the interaction between the population of GABA interneurons and the population of GABA efferent neurons (*c*_*ie*_). For low strength of interaction, GABAergic output increases as MePD excitation is increased, while excessively high interaction drives the activity of the population of GABA efferent (projection) neurons to a zero level. (**c**) Heat map depicting the combined mean output of glutamate and GABA projections. The dashed line represents the selected functional interaction strengths used in the model (see Table 1).

Another critical component of the system is the functional interaction strength between the populations of GABA interneurons and glutamate neurons, which facilitates oscillatory behaviour in the model by providing negative feedback between the two populations. Having fixed the kisspeptin excitation level to induce oscillatory dynamics (*K*_*p*_ = 2.3), we perform one parameter bifurcation analysis using the functional interaction strength between GABA interneurons and glutamate neurons (*c*_*il*_) as a bifurcation parameter. Our analysis reveals Hopf (**HB**), saddle node (**SN**) and homoclinic (**HC**) bifurcations that we then continue in two parameters (using the functional interaction strength between glutamate neurons and GABA interneurons (*c*_*li*_) as a second bifurcation parameter). This enables us to investigate the qualitative dynamics of the model under the variation of GABA-glutamate interaction (Figure 4(**a**)). The intersections of the Hopf and saddle node curves are associated with the location of Bogdanov-Takens (**BT**) points, which also gives rise to the homoclinic curve, while the location where two saddle node curves meet tangentially indicates the location of a co-dimension two cusp (**CP**) point. Our analysis confirms that the existence of oscillatory dynamics in the MePD neuronal network intricately depends on the balance between the inhibitory and excitatory interaction strengths. Specifically, insufficient or excessive strength of functional interaction between glutamatergic neurons and the population of GABA interneurons causes loss of oscillatory dynamics, while high inhibitory strength of interaction does not prevent oscillations, but rather makes the mean MePD projections’ output more inhibitory (Figure 4(**a**)).

**Figure 4.**
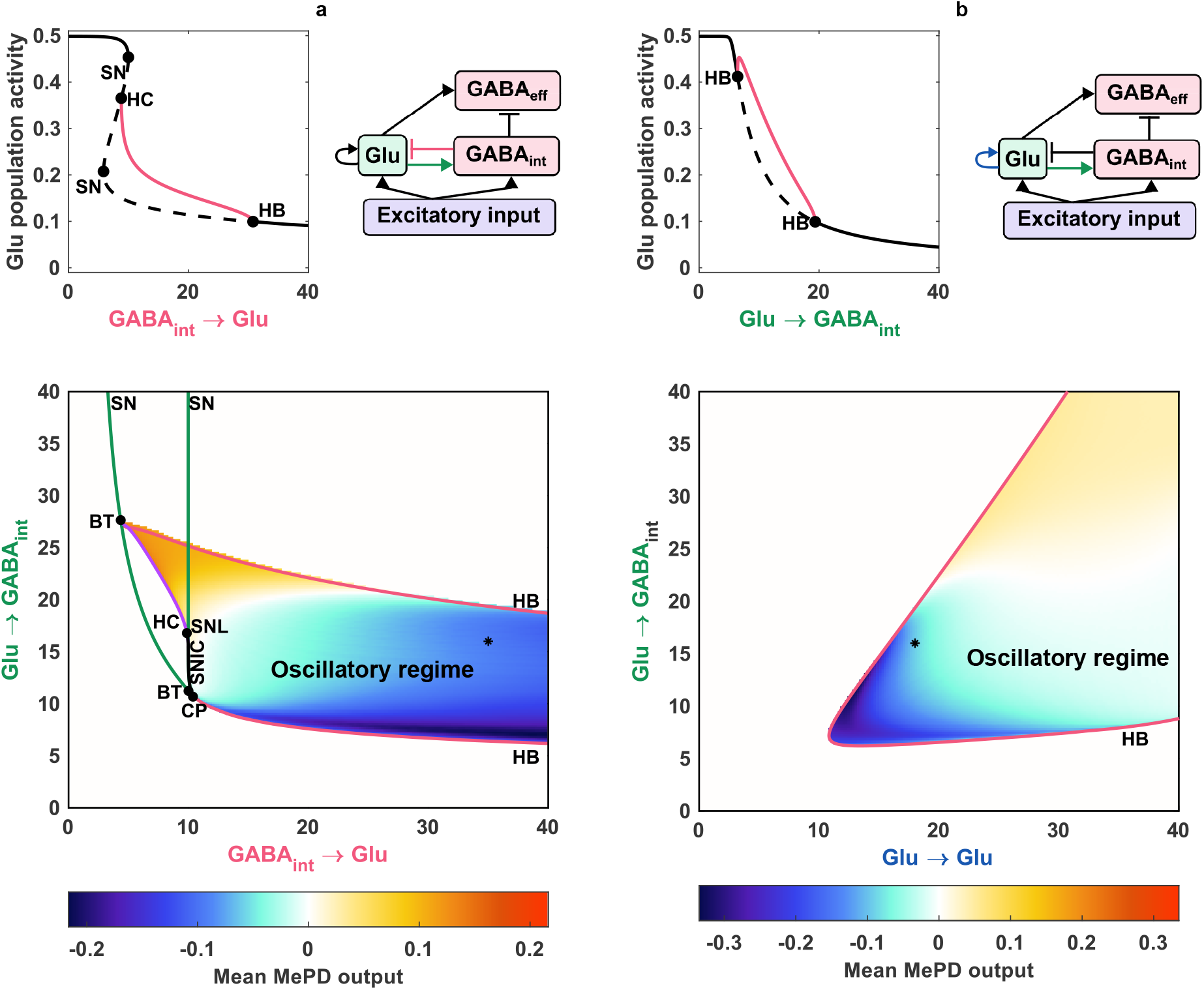
Effect of varying the MePD functional network connectivity on the MePD projections’ output dynamics. (**a**) Bifurcation analysis under varying functional interaction strength between the populations of GABAergic interneurons and glutamatergic neurons (*c*_*il*_ and *c*_*li*_). (**b**) Bifurcation analysis under varying strength of glutamatergic self-excitation (*c*_*ll*_) and functional interaction strength between glutamatergic neurons and GABAergic interneurons (*c*_*li*_). The network schematic diagrams highlight the connections under investigation. In the one-parameter bifurcation diagram, the red lines indicate the max of the limit cycle solutions branch, and circle markers denote Hopf (**HB**), saddle node (**SN**), and homoclinic (**HC**) bifurcation points, respectively. These points are used for the two-parameter continuation. In the two-parameter bifurcation diagrams, red, green, and purple lines represent Hopf (**HB**), saddle-node (**SN**) and homoclinic bifurcation curves (**HC**), respectively. The circular markers depict the Bogdanov-Takens (**BT**), saddle node loop (**SNL**) and cusp (**CP**) points. The heat map represents the mean MePD projections’ output in the oscillatory region. The star marker identifies the parameter values in Table 1.

Next, we consider the combined effects of the glutamatergic population’s self-excitation and its excitatory input to the population of GABA interneurons. We perform one-parameter bifurcation analysis using the strength of functional interaction between glutamate neurons and GABA interneurons (*c*_*li*_) as a bifurcation parameter, identifying Hopf (**HB**) bifurcation points (Figure 4(**b**)). Increasing the strength of functional interaction between glutamatergic population and GABA interneurons causes a decrease in glutamatergic population activity as the effect of the negative feedback loop is amplified, resulting in higher inhibitory interaction between GABA interneurons and the glutamatergic population. We then continue the detected (**HB**) points in two parameters (glutamatergic self-excitation strength (*c*_*ll*_) and strength of functional interaction between glutamate neurons and GABA interneurons (*c*_*li*_)) to define the oscillatory region in two-parameter space (Figure 4(**b**)). We find that higher levels of self-excitation require higher strength of functional interaction between glutamate neurons and GABA interneurons in order to give rise to oscillatory dynamics. Increasing the strength of glutamatergic functional interaction switches the mean MePD projections’ output from negative to positive.

The above analysis indicates that the intricate excitatory/inhibitory MePD projections’ balance is heavily dependent on the MePD circuit’s functional interactions. It also suggests that coordinated changes in the interaction strengths between the circuit’s neuronal populations may be a critical regulator of the MePD projections’ output, and thus its modulatory effect on the GnRH pulse generator.

### (d) MePD projections’ dynamic modulation of GnRH pulse generator activity

Having analysed the MePD GABA-glutamate neuronal network behaviour and the effects of different model parameters on the mean MePD projections’ output, we now investigate our coupled MePD-KNDy network model (see Coarse-grained Model of ARC KNDy population with MePD input for full model details). The aim here is to characterising the differential effects of MePD dynamic projections’ output on GnRH pulsatility. In essence, coupling the MePD and KNDy models results in feeding external periodic input from the MePD neuronal network with the KNDy neuronal network (a.k.a. GnRH pulse generator). As the timescales of the two network models are significantly different (MePD neuronal network evolves on a timescale of seconds while the KNDy neuronal network operates on a timescale of minutes), in the coupled model there is more than one frequency found in the periodic trajectory which now evolves on a torus, i.e. the limit cycle becomes a limit torus solution (Figure 5(**a**)). In the case when the extended system has two incommensurate frequencies, the trajectory is no longer closed, leading to quasi-periodic dynamics. This is not surprising as it is well established that in response to periodic input, relaxation oscillators such as the KNDy network model [6] can exhibit complex dynamics, like quasi-periodicity [38, 39].

**Figure 5.**
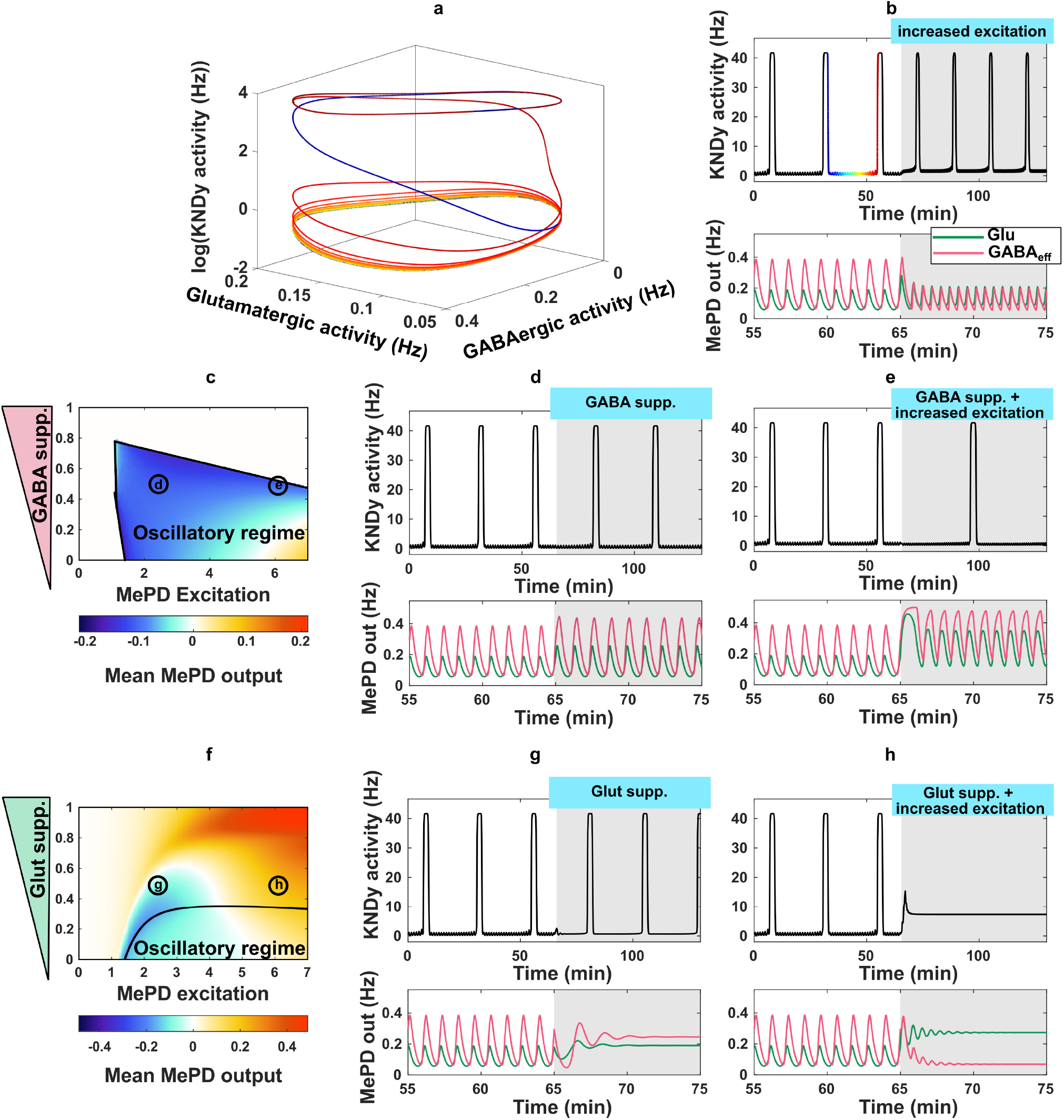
Differential effects of MePD projections on the dynamics of the GnRH pulse generator. (**a**) Providing oscillatory input to the KNDy network model leads to quasi-periodic behaviour evolving on a torus; we use a colour scheme to illustrate time evolution on this torus. (**b**) Increasing excitation in the MePD leads to a decrease in GABAergic tone, and as a result a decrease in interpulse interval (IPI) in the KNDy system. (**c**) Two-parameter bifurcation diagram (MePD excitation (*K*_*p*_) and GABAergic interaction suppression coefficient (*β*_1_)) showing the region where the MePD exhibits oscillatory dynamics and the mean MePD output superimposed on it. (**d**) Suppression of GABAergic interaction evenly increases both GABAergic and glutamatergic tone, leaving the overall MePD output as before the suppression, hence KNDy IPI remains the same. (**e**) Conjunction of the suppression of GABAergic interaction strength with increased MePD stimulation increases the proportion of GABAergic projections’ output, resulting in an increase in KNDy IPI. (**f**) Two-parameter bifurcation diagram (MePD excitation (*K*_*p*_) and glutamatergic interaction suppression coefficient (*β*_2_)) showing the region where the MePD exhibits oscillatory dynamics with superimposed mean MePD projections’ output. (**g**) Suppression of glutamatergic interaction strength leads to a loss of oscillatory dynamics in the MePD. Inhibitory GABAergic output is still higher than the excitatory glutamatergic output, leading to the return of periodicity in the KNDy system and leaving KNDy IPI as before the suppression. (**h**) Combining the suppression of glutamatergic interaction strength with the increased excitation of the MePD overstimulates ARC KNDy network, resulting in a transition to quiescent dynamics. A more detailed versions of the two-parameter bifurcation diagrams depicted in (**c**) and (**f**) are provided in Figure S2.

Next, we validate the coupled model by reproducing in vivo experiments where the effect of optogenetic stimulation of MePD kisspeptin neurons on LH pulse frequency was investigated [12]. To simulate the effects of optogenetic stimulation, we increase the kisspeptin level of excitation within the MePD, which results in activation of the GABA-GABA component of the MePD neuronal circuit, resulting in a decrease in the activity of the population of GABA efferent neurons’(Figure 5(**b**)). Consequently, the reduction of the inhibitory tone in the MePD projections’ output under increased excitation promotes a decrease in the interpulse interval (IPI) in the KNDy system (Figure 5(**b**)).

To investigate the effects of suppression of GABAergic interaction strength on the system’s dynamics, we compute a two-parameter bifurcation diagram for a range of MePD excitation (*K*_*p*_) and GABA functional interaction strength suppression coefficient *β*_1_ (see Figure 1(**b**)), the latter describing the strength of GABA receptor antagonism (Figure 5(**c**)). We find that complete suppression of GABAergic interaction leads to the loss of oscillatory dynamics in the MePD circuit via its effect on the negative feedback loop between the populations of GABA interneurons and glutamatergic neurons. It is common that the effect of pharmacological blockers is modelled by complete suppression of functional interactions [13], but in reality only partial suppression may occur. Here, we show that partial blocking of GABAergic interaction is sufficient to decrease the MePD projections’ output under increased excitation (Figure 5(**c**)). On the other hand, reduction of GABAergic functional interaction strength leads to a decrease in the inhibitory coupling between the population of GABA interneurons and the glutamatergic population and the population of GABA efferent neurons, hence increasing both glutamatergic and GABAergic tone in the MePD (Figure 5(**d**)). However, the difference between the inhibitory and excitatory tone remains relatively constant as before the suppression, resulting in an unperturbed KNDy network interpulse interval. Combining suppression of GABAergic interactions with increased excitation in the MePD also increases the activity in the populations of glutamatergic neurons and GABAergic efferent neurons, but also amplifies the difference between inhibitory and excitatory mean MePD projections’ output (Figure 5(**e**)). As a result, the MePD projections’ input to KNDy becomes more inhibitory, leading to an increase in the KNDy network interpulse interval.

Similarly, to mimic the effects of a glutamate receptor antagonist, we decrease the strength of glutamatergic interactions in the model and observe that oscillatory dynamics rapidly cease (Figure 5(**f**)). We also observe that under partial suppression of glutamatergic interaction strength and increased excitation, there is a transition from inhibitory to excitatory tone in the system, while complete abolition of glutamatergic interactions causes the mean MePD output to be exclusively excitatory for all levels of excitation (Figure 5(**f**)). To preserve inhibitory output from the MePD under low levels of excitation, we consider a partial suppression of the functional interaction strength associated with the population of glutamatergic neurons. For lower levels of excitation (still enabling oscillatory dynamics), moderate suppression of glutamatergic interaction strength results in a loss of oscillatory MePD dynamics, while keeping the mean MePD output relatively constant (and inhibitory); hence the extended system dynamics transitions from evolving on a limit torus to a limit cycle while the IPI of the KNDy system remains unaffected (Figure 5(**g**)). Suppression of glutamatergic functional interaction strength combined with an overall increase in excitation of the MePD network causes a switch from inhibitory to excitatory mean MePD output (due to the presence of GABA-GABA interactions). This results in over-stimulation of the KNDy network associated with a transition from a pulsatile to a quiescent mode of operation due to a depolarisation-block-type of phenomenon (Figure 5(**h**)).

### (e) How stimulation of MePD projections modulates GnRH pulse generator activity

The suppressive effect of MePD GABA projections’ output on the GnRH pulse generator frequency has been recently confirmed experimentally by optogenetically stimulating MePD GABAergic terminals in the ARC [16]. Here, using our coupled model, we interrogate the role of both GABAergic and glutamatergic projections of the MePD circuit in modulating KNDy dynamics. In the previous section the weights of GABA and glutamate projection output were set to be equal (*j*_*l*_ = *j*_*e*_ = 1). Here in order to account for direct optogenetic simulation of these projections as carried out in [16] we increase the weight of the respective projection output in the model (i.e. for simulating the experiments presented in [16] we increase the weight of GABA projections output in the model) and vary the stimulation level in the MePD circuit (*K*_*p*_). Specifically, if we set *j*_*e*_ = 1.5 and *j*_*l*_ = 0.5 we are able to show that increasing the stimulation level (*K*_*p*_) in the coupled system initially produces no change in the KNDy IPI, followed by an exponential increase in the KNDy inter pulse interval corresponding to cessation of the LH pulsatile dynamics (Figure 6(**a**)). These results conform with in vivo stimulation of MePD GABA projections, where a decrease in LH pulsatility (demonstrated by increase in inter pulse interval) eventually leading to loss of pulsatile dynamics have been observed at 10 and 20 Hz stimulation of GABA projections, respectively [16].

**Figure 6.**
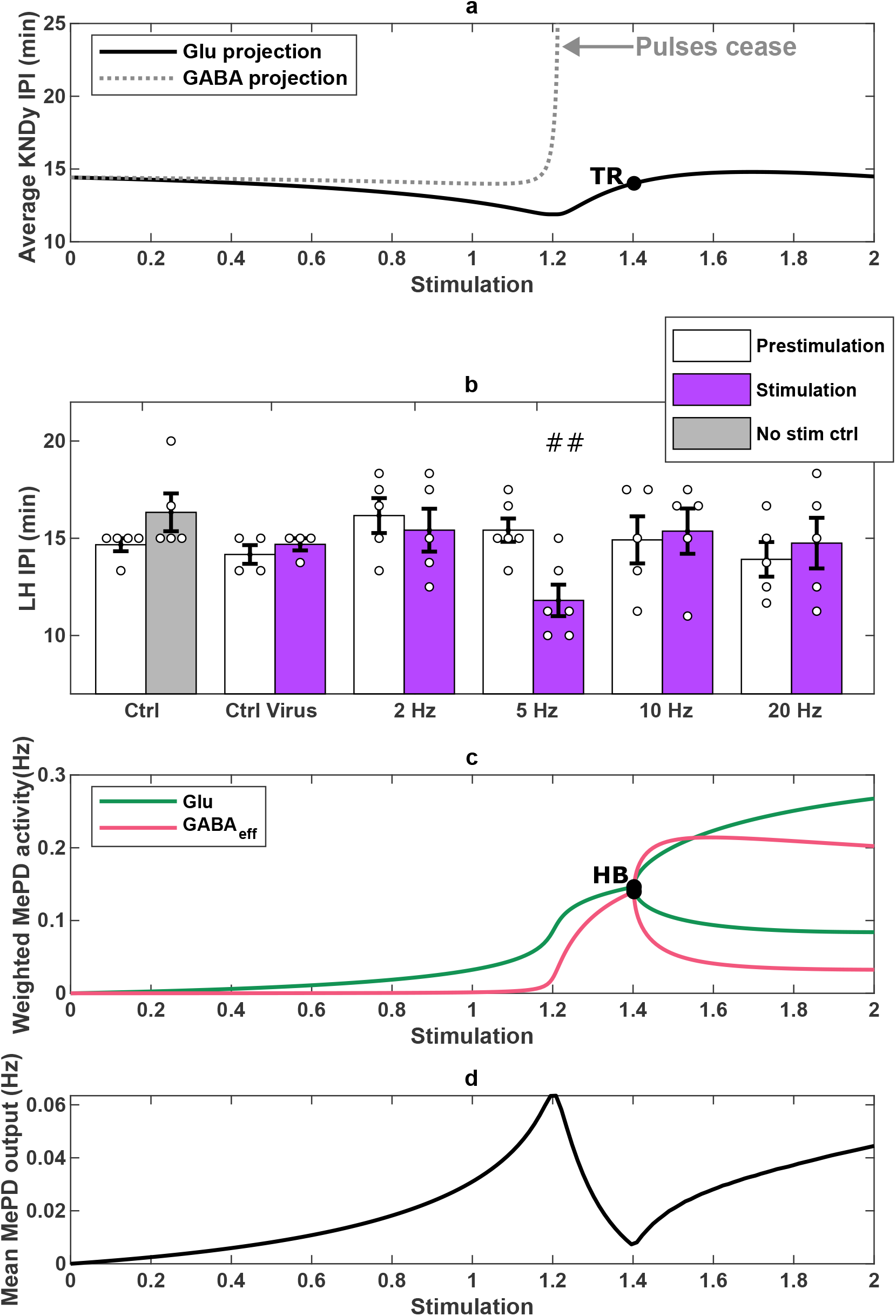
In silico simulation of MePD projection stimulation and non-monotonic LH interpulse interval response to stimulation of MePD glutamatergic projection in vivo. (**a**) Simulation of GnRH pulse generator response to stimulation of GABA and glutamate MePD projections. Mimicking the effects of GABA projection stimulation leads to the loss of oscillatory dynamics in the KNDy activity as stimulation level is increased. During glutamate projection stimulation in silico KNDy IPI initially declines and then rises back as the level of stimulation (*K*_*p*_) (given in arbitrary units (a.u.)) increases. A torus bifurcation (**TR**) denotes the transition from limit cycle to limit torus in the extended MePD-KNDy network model. The location of **TR** and **HB** align. (**a**) Summary of average LH interpulse interval (IPI) for non-stimulation control (*n* = 5), control virus (*n* = 4) and optical stimulation at 2 Hz (*n* = 5), 5 Hz (*n* = 6), 10 Hz (*n* = 5), and 20 Hz (*n* = 5) of MePD glutamatergic projection terminals in the ARC. We depict the mean *±* SEM, and individual data points for each animal as circles in the histogram plots. Stimulation at 5 Hz results in a statistically significant suppression of LH pulse frequency ^##^*F* = 12.8517, *p* = 0.0050 vs control period for 5 Hz optical stimulation group. (**c**) The external MePD projections’ input to the KNDy network, showing glutamatergic and GABAergic contributions as a function of the level of stimulation (in a.u.). Hopf point (**HB**) point denotes the location of the transition from stationary to oscillatory dynamics in the MePD circuit. (**d**) The variation in the weighted mean MePD output under different levels of stimulation.

Now, we employ the same strategy to simulating the effects of glutamate projections’ stimulation (by fixing *j*_*l*_ = 1.5 and *j*_*e*_ = 0.5). Our model simulations show that under increasing excitation of the MePD network, the inter pulse interval of KNDy population activity initially decreases before increasing and then returning to its initial IPI (Figure 6(**a**)). Furthermore, bifurcation analysis of the extended model demonstrates that the system undergoes a torus bifurcation (**TR**), associated with the switch from limit cycle dynamics to dynamics evolving on a limit torus, which we depict in (Figure 5(**a**)). The observed non-monotonic behaviour of the coupled system in this case is counterintuitive, given the excitatory role of glutamate. Nevertheless, selective in-vivo optogenetic stimulation of the MePD glutamatergic projections in the ARC with increasing levels of stimulation confirms our model predictions as shown in (Figure 6(**b**)). As expected, given the excitatory nature of the glutamatergic projections, sustained stimulation at 5 Hz results in a significant decrease in LH interpulse interval from 15.42 ± 0.60 min to 11.81 ± 0.81 min. However, further increase in the frequency of stimulation (at 10 Hz and 20 Hz) restores the pre-stimulation IPI levels, as predicted by our modelling.

Moreover, our modelling allows us to explore potential mechanisms that govern the non-monotonic response in the GnRH pulse generator found experimentally as described above (Figure 6(**c-d**)). Continuation analysis of the MePD model dynamics indicates that the decrease in the KNDy network IPI observed in the model is due to the amplification of glutamatergic activity in the MePD network, while GABAergic tone remains very close to zero (see Figure 6(**c**) at stimulation level ≈ 1.2). Further increase in MePD network excitation, however, switches the balance in excitatory/inhibitory MePD projections’ output (i.e. MePD input to KNDy)(Figure 6(**d**), which in turn promotes an increase in the KNDy IPI. The MePD network model undergoes a Hopf bifurcation (**HB**), the location of which is associated with the location of the torus bifurcation (**TR**) in the extended MePD-KNDy network model (Figure 5(**a-b**)), demonstrating that the transition from limit cycle to limit torus occurs due to a change in the qualitative dynamics of the MePD network model.

## 4. Discussion

In our study, we have introduced and systematically investigated a model incorporating the interplay between GABA and glutamate neuronal populations within the MePD. This model was coupled to a GnRH pulse generator model [6, 7], allowing us to validate it against experimental findings from [12, 13] as well as offering insights into how perturbations in the MePD could impact the activity of the GnRH pulse generator. Our model could serve as a versatile tool for investigating broader MePD circuit effects, such as for example those stemming from the interactions between urocortin and the GABA/glutamate neuronal populations [40]. The utility of our modelling approach lies in its ability to interrogate the neuronal mechanisms that enable the MePD to modulate the dynamics of the GnRH pulse generator, hence enabling us to better understand the effects of environmental and psychosocial factors on reproductive function, such as pubertal timing [9–11] and modulation of LH secretion [12, 13].

A model of the MePD circuit coupled to the GnRH pulse generator has been previously studied under stationary MePD circuit dynamics [13]. In this mode, the system functions like a collection of independent neuronal oscillators characterised by a constant (averaged) population level of activity. However, it is important to note that while this study has offered valuable insights, the actual patterns of MePD activity are likely to be more complex. Indeed, [31] shows changes in the MePD neuronal network’s oscillatory activity in vivo associated with sex-specific differences in the encoding of social stimuli and sexual experience. Here, we demonstrate that the extended model is able to reproduce experimental findings [12, 13], suggesting the plausibility of an oscillatory mode of MePD circuit activity. In fact, rhythmicity of neuronal populations is a characteristic feature of neuronal synchronisation, allowing the neuronal networks to manage and process complex stimuli [41]. However, to confirm or reject the hypothesis about the importance of oscillations in the MePD neuronal networks, further experiments involving recordings of calcium activity in individual GABA and/or glutamate neurons in the MePD, and how they synchronise, would be required.

Under oscillatory MePD circuit behaviour, our modelling shows that an increase in the excitatory input to the MePD system decreases GABAergic MePD output due to the activation of GABA-GABA disinhibition, while glutamatergic output remains consistent. This finding indicates that GABAergic MePD output is sensitive to stimulatory inputs, while glutamatergic output is likely to play more of a balancing role. This is consistent with the established role of amygdala in reproductive function modulation, as lesions to the MePD have been shown to advance puberty [42] and prevent stress-induced suppression of LH pulses [43]. On the other hand, optogenetic stimulation of kisspeptin neurons in the MePD [12] increases LH pulse frequency, and administration of peripheral kisspeptin inhibits neuronal activation in the amygdala as well as increases LH secretion [44], which can be explained by a decrease in the activity of GABAergic efferent neurons.

Here, we have modelled the effects of pharmacological interventions via partial suppression of MePD circuit functional interactions and studied how different levels of suppression affect qualitative dynamics in the model; this is in contrast to our previous work [13] where we assumed complete suppression of signalling. In reality, however, complete suppression is unlikely to be the case, as neuronal cells may respond to pharmacological interventions by upregulating receptors or modifying their signalling pathways to compensate for the inhibited receptors. When modelling partial GABA signalling suppression, the inhibitory component of MePD output increases because there is not enough GABA-GABA disinhibition, but at the same time glutamatergic activity also goes up due to decreased inhibition from the GABAergic population, balancing out the inhibitory output. This compensation mechanism could provide an alternative explanation as to why solely GABA receptor antagonism does not change LH pulsatility [13].

Feeding oscillatory MePD projections’ output into the KNDy network allowed us to consider dynamic upstream modulation of the KNDy network rather than a constant input as considered in our previous work [7, 13]. We investigated how such dynamic input changes the response of the KNDy relaxation oscillator. Specifically, we have demonstrated that when the KNDy relaxation oscillator receives a dynamic input of a significantly different frequency, this can result in a complex quasi-periodic pulse pattern, i.e. irregular-shaped pulses with no change in interpulse interval. Quasi-periodicity has also been identified in other relaxation oscillator systems subject to periodic inputs, indicating that systems characterised by quasi-periodic behaviour often possess a level of resilience against external perturbations [38, 39]. On the other hand, the transition from oscillatory to constant input, which we observe during the suppression of glutamatergic functional interactions, makes the KNDy system far more sensitive to the magnitude of the change. Previous modelling work suggests high sensitivity of the KNDy network to the magnitude of constant external stimuli that can lead to cessation of GnRH pulses [7]. However, to maintain a functional reproductive system, the GnRH pulse generator must be resilient to small perturbations that can arise from changes in the MePD (or other upstream brain regions) output due to environmental stimuli, which is more plausible under periodic input modulation.

Previously published data identified that stimulation of MePD GABA projections in the ARC modulates GnRH pulse generator activity [16]. Using our extended mathematical model we interrogated the role of the glutamatergic projections in such modulation, which has not been shown previously. The model analysis predicted a possible non-monotonic response of KNDy activity. These model predictions were experimentally confirmed in vivo, suggesting a novel excitatory MePD glutamatergic projection capable of impacting the KNDy network, and hence modulating the GnRH pulse generator. Specifically, we show that stimulation at 5 Hz leads to a statistically significant decrease in LH interpulse interval, but further increase in stimulation frequency (10 Hz and 20 Hz) produces no significant change. Considering the excitatory function of glutamate, one might question why an increase in stimulation does not lead to a further decrease in LH IPI or even the complete loss of pulsatility due to potential overstimulation (or in other words depolarisation block). Using our extended mathematical model we reproduced this non-monotonic stimulus-response relationship, providing insight into potential mechanisms though which glutamatergic projections modulate GnRH pulse generator activity. In the model, as the excitatory drive increases, so too does the response of excitatory projection neurons, leading to positive correlation between excitatory input and neuronal response, similar to the effect observed during 5 Hz stimulation of glutamatergic projections. However, when excitation becomes stronger, which mimics the effects of the higher frequency optogenetic stimulation, the system reaches a point where further increase in the excitatory drive does not lead to a proportional increase in the neuronal response. Excessive stimulation may also trigger compensatory mechanisms that counteract the increased activity, leading to a limited net change [45]. As we observe in the model, the inhibitory projections could become engaged to maintain the balance, counteracting the excitatory drive and contributing to a non-monotonic response. Based on the model analysis, we argue that balanced feed-forward excitation and feed-forward inhibition ensure that the overall excitability of the KNDy network is robust to the elevated excitatory input, permitting GABAergic projections to exert ‘inhibitory brake’, thus constraining the LH IPI to pre-stimulation levels.

Here we propose a mathematical model and explore the influence of MePD activity on GnRH pulse generator dynamics. The modelling and analysis approach we have used could be applied to gain insight into the behaviour of other brain regions involved in modulation of the GnRH pulse generator. Additionally, given the phenomenological nature of our MePD model, in the future, the extended MePD-KNDy model could be used to interrogate the effects of other stimulatory neuronal populations signalling to the MePD GABA-glutamate circuit. This could then be employed to perform in silico simulations to interpret and/or predict experimentally observed effects on the GnRH pulse generator and reproductive function.

## Appendix

### Data Accessibility

The code to reproduce the analysis and data can be found in [GitHub repository](https://github.com/ktrnnchprnk/MePDmodel).

### Authors’ Contributions

Kateryna Nechyporenko, Conceptualisation, Investigation, Methodology, Software, Writing – original draft; Margaritis Voliotis, Conceptualisation, Investigation, Writing – review and editing; Xiao Feng Li, Conceptualisation, Data curation, Investigation; Owen Hollings, Conceptualisation, Data curation, Investigation; Deyana Ivanova, Conceptualisation, Investigation; Jamie Walker, Conceptualisation, Investigation, Writing - review and editing; Kevin O’Byrne, Conceptualisation, Investigation, Writing - review and editing; Krasimira Tsaneva-Atanasova, Conceptualisation, Investigation, Writing - review and editing.

### Funding

KTA gratefully acknowledges the financial support of the EPSRC via grant EP/T017856/1. KTA and MV gratefully acknowledge the financial support of the BBSRC via grant BB/W005883/1. KOB and XFLI gratefully acknowledge the financial support of the BBSRC via grant BB/W005913/1. JJW gratefully acknowledges the financial support of the MRC via grants MR/N008936/1 and MR/T032480/1.

### Disclaimer

For the purpose of open access, the author has applied a ‘Creative Commons Attribution (CC BY) licence to any Author Accepted Manuscript version arising.

---

**Figure S1.**
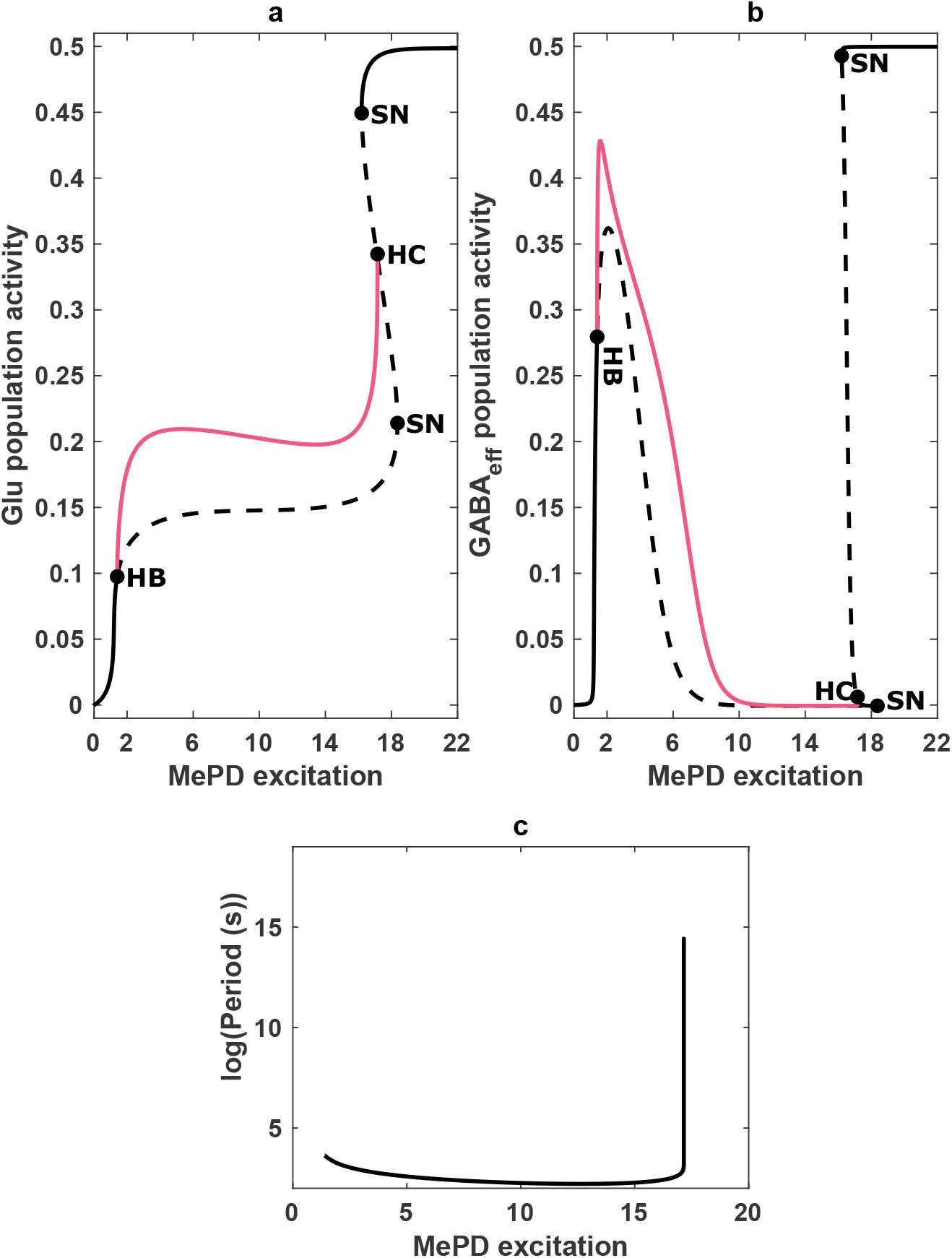
Extended bifurcation diagrams of kisspeptin-mediated excitatiry input to GABA-glutamate circuit from Figure 2 for (**a**) *G*_*l*_ and (**b**) *G*_*e*_ dynamics. Circle markers denote Hopf (HB), homoclinic (HC) and saddle node (SN) bifurcations. The red line indicates the maximum amplitude of the limit cycle solutions branch. (**c**) Excitatory input vs. log Period of oscillations. The period, firstly, decreases, followed by a rapid increase, signifying homoclinic bifurcation.

**Figure S2.**
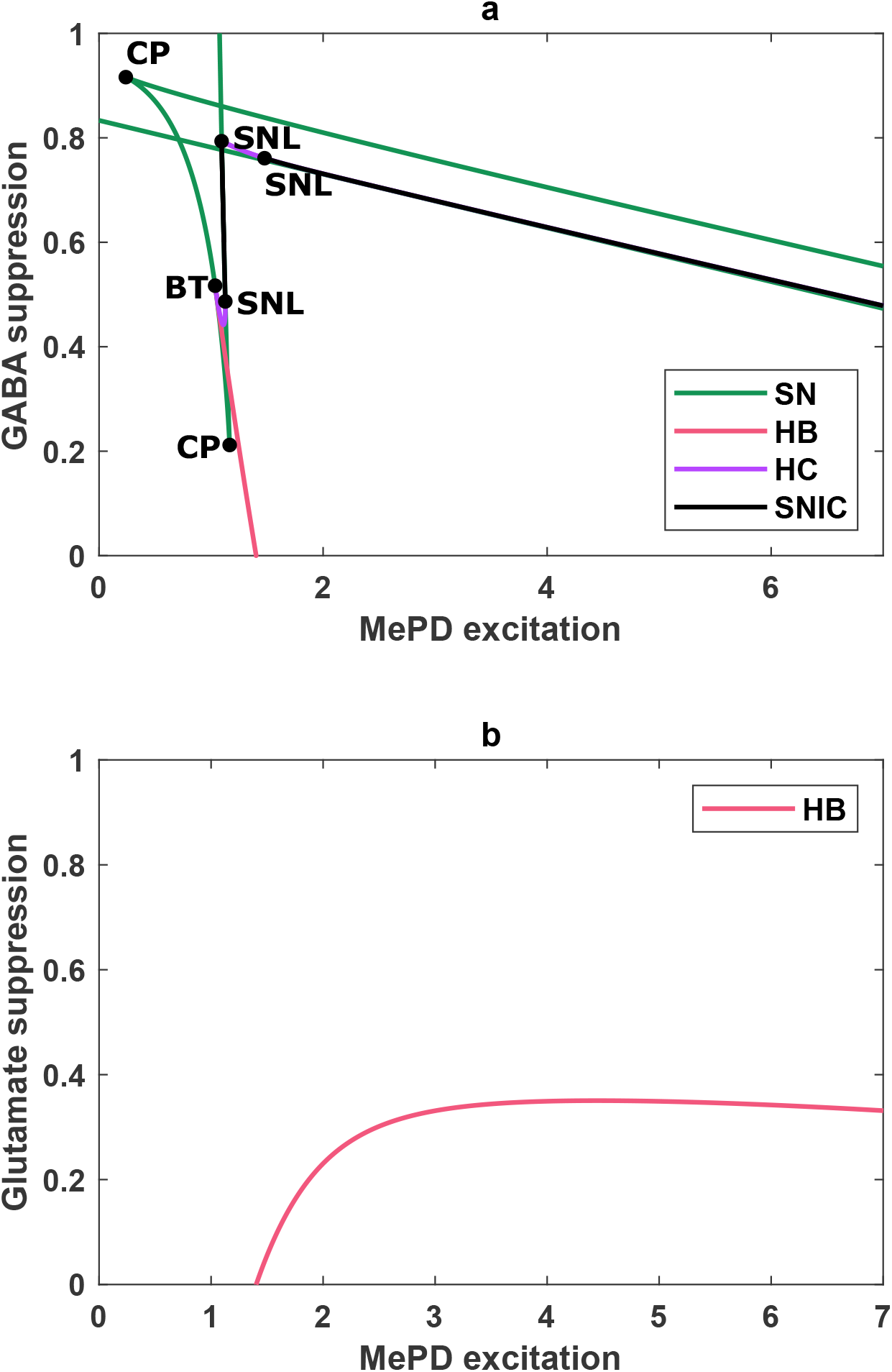
Two parameter bifurcation diagrams, used to identify the oscillatory regions in Figure 5(**c,f**). (**a**) Two-parameter bifurcation diagram of MePD excitation (*K*_*p*_) and GABAergic interaction suppression coefficient (*β*_1_). (**b**) Two-parameter bifurcation diagram of MePD excitation (*K*_*p*_) and glutamtergic interaction suppression coefficient (*β*_2_). Red, green, purple lines represent Hopf (**HB**), saddle-node (**SN**) and homoclinic bifurcation curves (**HC**), respectively. The circular markers depict Bogdanov-Takens (**BT**), saddle-node loop (**SNL**) and cusp (**CP**) points.

